# Evaluation of fully oxidized β-carotene as a feed ingredient to reduce bacterial infection and somatic cell counts in cows with subclinical mastitis

**DOI:** 10.1101/2020.10.12.335463

**Authors:** S McDougall

**Author notes:** **Abbreviations** CNS: coagulase negative staphylococci; DIM: days in milk; OxBC: oxidised β-carotene; SCC: Somatic cell counts; TLR: Toll-like receptors.

## Abstract

**Aims:** To assess the effect of oral supplementation with fully oxidised β-carotene (OxBC) on bacteriological cure, incidence of clinical mastitis, and somatic cell counts (SCC) in cows with subclinical intramammary infection.

**Methods:** Cows from four dairy herds were enrolled in early lactation if they had quarter-level SCC >200,000 cells/mL and they had a recognised bacterial intramammary pathogen in one or more quarters. They were randomly assigned to be individually fed from Day 0, for an average of 40 days, with 0.5 kg of a cereal-based supplementary feed that either contained 300 mg of OxBC (treatment; n=129 quarters) or did not (control; n=135 quarters). Quarter-milk samples were collected on Days 21 and 42 for microbiology and SCC assessment. Bacteriological cure was defined as having occurred when the bacteria present on Day 0 were not isolated from samples collected on Days 21 or 42. Clinical mastitis was diagnosed and recorded by herdowners up to Day 42.

**Results:** The bacteriological cure rate was greater for quarters from cows in the treatment group (13.9 (95% CI=4.1–23.7)%) than for quarters from cows in the control group (6.9 (95% CI=4.8–9.1)%; p=0.02). The prevalence of quarters that were infected on Day 42 was less in cows in the treatment group (79.9 (95% CI=62.3-97.6)%) than the control group (88.2 (95% CI=78.4-97.9)%; p=0.009). The incidence of quarters diagnosed with clinical mastitis by Day 42 was lower in cows from the treatment group (1/129 (0.78 (95% CI=0.02-4.24)%) than in cows from the control group (6/135 (4.44 (95% CI=1.65-9.42)%; p=0.03). Mean quarter-level SCC did not differ between treatment groups (p=0.34).

**Conclusions and Clinical Relevance:** Feeding 300 mg/cow/day of fully oxidised β-carotene resulted in a higher bacteriological cure rate, a lower prevalence of intramammary infection following 6 weeks of feeding, and a lower incidence of clinical mastitis compared to untreated controls. This offers a non-antimicrobial approach to reducing prevalence of intramammary infection in dairy cows.

## Introduction

Intramammary infection of dairy cows is common and results in increased somatic cell counts (SCC) at quarter, cow and bulk tank milk level, increased risk of clinical mastitis, and of culling with resultant negative economic impacts on dairy farms (Halasa *et al.* 2007). Studies in Australia and New Zealand have assessed total lactation milk production losses associated with subclinical intramammary infection to be between 14% and 23% (Morris 1973; Mwakipesile *et al.* 1983). A non-peer reviewed New Zealand study found that antimicrobial treatment of cows with elevated SCC resulted in an increased cure rate of 10-20% compared with non-treated quarters (Douglas *et al.* 1997). The cost effectiveness of therapy of subclinical mastitis during lactation is dependent on the relative cure rates of treated versus untreated quarters, rate of transmission of infection within the herd, retention pay-off for culling, and the costs of treatment, and is negative in some situations (Swinkels *et al.* 2005). Additionally, with increasing concern about antimicrobial use, treatment of subclinical mastitis may be seen as inappropriate antimicrobial stewardship.

Studies from the 1980s reported a positive relationship between udder health and concentrations of vitamin A and beta-carotene in plasma of postpartum dairy cows (Chew *et al.* 1982; Johnston and Chew 1984). More recently, oral supplementation of peripartum dairy cows fed a total mixed ration with β-carotene tended to reduce the proportion of multiparous cows with SCC >200,000 cells/mL (Oliveira *et al.* 2015), and hyper-supplementation with oral vitamin A resulted in reduced SCC after 60 days (Jin *et al.* 2014). Both vitamin A and β-carotene affect immune function and vitamin A deficiency is associated with immunosuppression (Scherf *et al.* 1994; Sordillo *et al.* 1997). Vitamin A improves the antioxidant defence systems against oxidative stress (Chew and Park 2004). An inverse association exists between the Rapid Mastitis Test score and milk concentrations of vitamin A and beta-carotene (Chew *et al.* 1982). However, in another study, no clear effect of supplementation with vitamin A or beta-carotene pre-dry off on polymorph neutrophil (PMN) function in milk could be demonstrated (Tjoelker *et al.* 1990). Oral supplementation with 1.2 g/cow/day of beta-carotene in the peripartum period tended to reduce the proportion of cows with a SCC >200,000 cells/ml and to reduce the risk of retained fetal membranes in multiparous cows (Oliveira *et al.* 2015). β-carotene may act independently of vitamin A as an oxygen radical scavenger, enhancing innate immunity by upregulation of toll like receptors (TLR), and modulating cytokine concentrations (Burton *et al.* 2014).

Recently a product has been developed containing a novel combination of β-carotene-oxygen copolymers formed by full oxidation of β-carotene. This product does not possess the ability to activate the vitamin A receptor system nor does it possess the antioxidant activity that has been purported as a mode of action of intact β-carotene (Burton *et al.* 2014; Johnston *et al.* 2014). *In vitro* studies demonstrated that fully oxidised β-carotene (OxBC) increased plasma membrane content of TLR subtypes 2 and 4 and their co-factor CD14 in several cultured cell types, including monocytes, fibroblasts, and endothelial cells. Additionally, pre-treatment with OxBC potentiated the lipopolysaccharide-induced increase in concentrations of some cytokines and increased phagocytic activity of monocytes (Johnston *et al.* 2014).

The immunomodulatory actions of OxBC have been evaluated in a dairy calf model of bovine respiratory disease. This disease is characterised by an excessive, neutrophil-driven, inflammatory response to the pathogen, which leads to pulmonary pathology and potentially death. The study demonstrated that, in *Mannheimia haemolytica* challenged calves, oral supplementation with OxBC reduced pulmonary inflammation via neutrophil-apoptosis and subsequent removal of apoptotic cells. Importantly, supplementation with OxBC was not associated with a reduction in circulating leukocytes, and supporting *in vitro* studies showed that OxBC did not inhibit neutrophil function (Duquette *et al.* 2014).

Oxidised β-carotene potentially has a role in managing bovine mastitis via upregulation of TLR-4 and CD-14, resulting in earlier recognition and removal of pathogens (De Schepper *et al.* 2008). Additionally, as demonstrated in the bovine respiratory disease model, OxBC may reduce the extent of neutrophil infiltration and hence reduce SCC in milk. Thus, OxBC potentially offers an non-antimicrobial approach to reducing the prevalence of intramammary infection and SCC.

We hypothesised that oral supplementation with OxBC of cows with subclinical intramammary infections would result in an increased bacteriological cure rate, a reduced incidence of clinical mastitis, and a reduced SCC relative to untreated controls.

## Materials and methods

### Animals and treatments

This study was undertaken with the prior approval of the Ruakura Animal Ethics Committee (AgResearch, Hamilton, NZ).

Cows from four spring calving dairy herds in the Waikato region of New Zealand were enrolled between August and November 2019. The herds were selected on the basis of maintaining good animal health records, undertaking routine herd testing, and agreement to follow the study protocol. Herd sizes varied from 424 to 799 lactating cows (Table 1). The primary feed during lactation for all herds was pasture containing ryegrass (*Lolium perenne*) and white clover (*Trifolium repens*), but all herds also supplemented cows with other feed types, as shown in Table 1. Cows were milked twice daily.

**Table 1.**
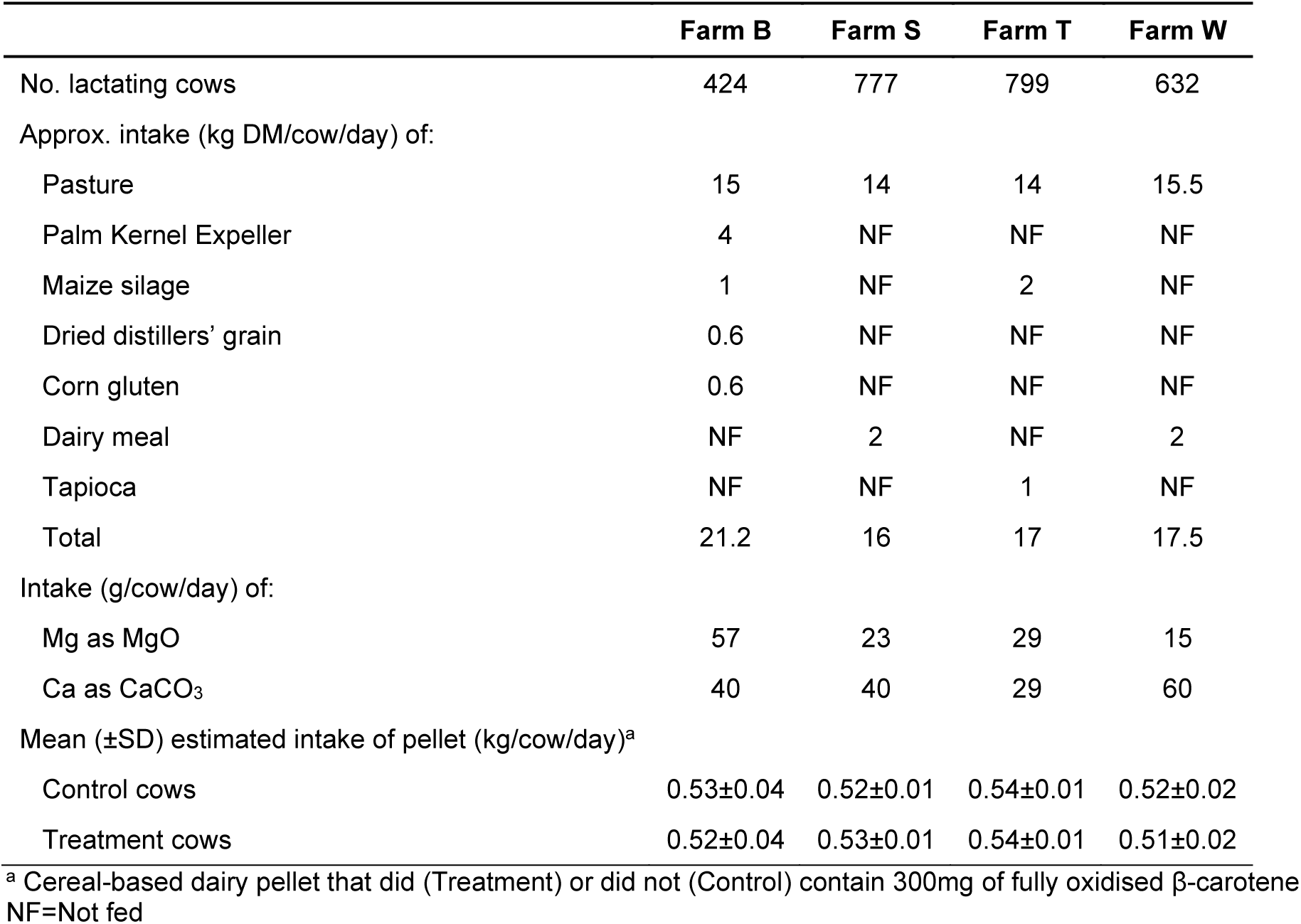
Summary of estimated dry matter (DM) intake of pasture and supplementary feeds, Mg and Ca, by lactating cows in four dairy herds, and intake of pelleted feed by cows that were supplemented with oxidised β-carotene (treatment) or not (control).

Herd testing was carried out on each farm between 6 and 15 days prior to the start of the study. Cows of any age were selected from each herd on the basis of having SCC >200,000 cells/mL at the herd test. Lists of cows to be drafted for milk sampling for the study were provided to the herd owner or manager. On the day of sampling trained research technicians assessed the teat end score (Mein *et al.* 2001) of selected cows and those cows with severe teat end damage (very rough) were excluded. Additionally, electronic, and on-farm records of treatments with antimicrobial and nonsteroidal anti-inflammatory drugs were assessed, and any cow treated in the preceding 14 days was excluded.

Two milk samples (~ 5 mL and ~ 25 mL) were collected from each quarter of each selected cow, following aseptic teat end preparation. These were held at 4°C and processed for microbiology within 24 hours of collection and submitted for quarter-level SCC determination within 72 hours of collection, as described below. Cows were then enrolled on the basis that they had a recognised bacterial intramammary pathogen in one or more quarters and quarter-level SCC >200,000 cells/mL.

Within each herd, enrolled cows were blocked by age (2 vs. >2 years), ranked on SCC recorded in the herd test, and then assigned randomly within sequential pairs of cows to control or treatment groups. All enrolled cows received 0.5 kg/day of a cereal-based pelletised feed that was formulated and mixed by a commercial feed mill (SealesWinslow, Morrinsville, NZ). The pellet contained 25% wheat, 25% maize, 30% palm kernel expeller, 8% broll, 4% dried distillers’ grain, 8% molasses, and 0.02% of a sweetener (rumasweet). The treatment supplement additionally included 0.6% of OxBC (OxC-beta Livestock, Avivagen, Ottawa, Canada), that is, 300 mg/cow/day of oxidised β-carotene.

Commencing on Day 0, within 6-12 days of initial milk sampling, the cereal pellet was fed to individual cows by research technicians at daily visits to each farm during milking, for an average of 40 (SD 2.2; min 37, max 42) days. The feed supplement was delivered using a measuring jug attached to a pole which allowed the technicians to stand behind the cows in the milking parlour and deliver the appropriate feed supplement to troughs located in front of the cows. To facilitate identification of study animals, cows were marked with paint of one of two different colours corresponding to the control or treatment groups. The delivery of feed supplement to each enrolled cow each day was recorded, as well as whether the feed supplement was fully consumed by each cow. The mass of feed supplement delivered each day to each group within each herd was calculated by weighing the feed supplement bags before and after each day’s feeding and recording the number of cows fed. The estimated daily offering of the feed supplement to cows on each farm is presented in Table 1.

Herd owners assessed cows daily and those with grossly evident signs of clinical mastitis were treated and recorded following normal farm protocol. Herd testing was performed again on each farm on Days 36, 55, 31, and 57 for Herds B, W, T, and S, respectively. Milk samples were also collected, as described above, from enrolled quarters on Days 21 and 42 for bacteriology and SCC.

### Laboratory procedures

Microbiology was undertaken at the Cognosco Laboratory (Morrinsville, NZ), following the procedures recommended by the National Mastitis Council, USA (Hogan *et al.* 1999). Briefly, 10 ml of milk was streaked onto a quarter of a 5% blood agar plate containing 0.1% esculin (Fort Richard, Auckland, NZ), and incubated at 37°C for 48 hours. The genus of bacteria was determined on the basis of colony morphology, Gram stain, and catalase reaction. Gram positive, catalase positive isolates were tested with the coagulase test to differentiate coagulase negative staphylococci (CNS) from *Staphylococcus aureus*. Gram positive, catalase negative were assessed by esculin reaction, CAMP test and growth in inulin and SF broth. Coliforms were sub-cultured on MacConkeys agar, triple iron sugar, citrate, and motility tests were performed. Where the identity of the isolate was unclear using conventional biochemical tests, isolates were submitted for matrix-assisted laser ionization time-of-flight (MALDI TOF) mass spectrometry (Pathlab, Tauranga, NZ). Where two distinct bacterial species were isolated from a milk sample these were reported individually and used in the analysis of spontaneous resolution or new intramammary infections. However, for reporting, they are coded as mixed major infections when a major pathogen (i.e. *S. aureus, Streptococcus uberis, Streptococcus dysgalactiae*) was isolated in conjunction with a minor pathogen (e.g. CNS or *Corynebacterium* spp.), or as mixed minor infections when two different minor pathogens were present. Samples were defined as contaminated if more than 2 distinct colony types were present.

SCC were determined using a fluoro-optic methodology (Foss, Hillerod, Denmark) at the laboratories of LIC (Riverlea, Hamilton, NZ). Results were forwarded as comma separated variable files which were loaded into a purpose-built database.

### Statistical analyses

The key outcome variable was resolution of infection (bacteriological cure). Bacteriological cure was defined as having occurred when the bacterial species present on Day 0 was not isolated from samples collected on either of Day 21 or 42. Cure was defined as occurring even when a different bacterial species was identified in samples collected on Days 21 or 42. Enrolled quarters that were diagnosed with clinical mastitis before Day 42 were defined as cure failures. Milk samples that were contaminated (n=3) or were not collected as the cow was not presented (n=1), were coded as a null.

Quarters defined as having developed a new intramammary infection where a bacterial species was isolated either at Day 21 or 42 that differed from that isolated on Day 0. Additional outcome variables were the prevalence of quarters that were infected on Day 42, quarter-level SCC on Days 0, 21 and 42, the proportion of quarters with SCC <200,000 cells/mL on Days 21 and 42, and cumulative incidence of clinical mastitis between Days 0 and 42. The quarter- and cow-level SCC, milk yield, and cumulative clinical mastitis incidence were also analysed.

The distribution of cows within herds, age groups (primiparous or multiparous), breed (≥12/16^th^ Friesian *vs.* other), and by days in milk (DIM) on Day 0 (categorised as 15–60, 61– 75, 76–90, ≥91) were compared between treatment groups using χ^2^ analysis. Cow-level SCC at the herd test preceding commencement of the study were natural log (ln) transformed and compared between treatment groups using one-way ANOVA.

The effect of treatment group on bacteriological cure, new intramammary infection, prevalence of infected quarters on Day 42 and cumulative incidence of clinical mastitis, was examined using multivariable models. Additional explanatory that were also considered during the modelling process were age group (primiparous vs. multiparous), breed (Friesian *vs*. other), quarter-level SCC on Day 0, quarter location (rear *vs*. fore), DIM at Day 0, and intramammary infection on Day 0 (categorised as major or minor). Initially bivariate analysis was undertaken using χ^2^ analysis for categorical variables and logistic regression analysis for continuous variables. Those variables associated (p<0.2) were then used in a manual forward stepwise model building process. Variables remained in the final models where p<0.05 or removal resulted in >20% change in the coefficient for the effect of treatment group. The final models were binary logistic regression models which included herd as a random effect and using robust variance estimates by specifying a variance-covariance matrix allowing for correlations of quarters and cows within herds. Attempts to create a model that included quarter within cow within herd resulted in a failure of the model to converge, due to the relatively small number of cows with multiple quarters enrolled. In addition, as no primiparous cows met the enrolment criteria in Herd S, models that included age also failed to converge and age was removed from subsequent models. Model fit was assessed using the Hosmer Lemeshow test, the link test and by assessing change in the Bayesian Information Criterion (BIC) when including or excluding variables.

Quarter-level SCC on Days 0, 21 and 42 were natural log (ln) transformed prior to multilevel repeated-measures generalised linear regression modelling with quarter nested within cow nested within herd. Fixed effects included treatment group and day (i.e. Days 0, 21 and 42). The interaction of treatment by day was forced into the model. Estimated marginal means and 95% CI were derived from the final model.

The proportions of quarters with SCC ≤200,000 cells/mL on Days 21 and 42 were analysed using a multilevel, repeated measures logistic regression model, with quarter nested within cow nested within herd. Fixed effects included treatment group and day (i.e. Days 21 and 42). Note by design only quarters with an SCC >200,000 cells/mL on Day 0 were included in the study, so there were no quarters with SCC ≤200,000 cells/mL on Day 0. The interaction of treatment by day was forced into the model. Estimated marginal means and 95% CI were derived from the final model.

The cow composite (herd test) SCC, milk yield (kg/cow/d) and milk solids (i.e. sum of kg of milk fat and milk protein/cow/day) from immediately prior to initiation of treatment and the next herd test during or just after the period of treatment were analysed using multilevel repeated-measures generalised linear regression modelling with cow nested within herd. SCC was natural log transformed for analysis. Fixed effects included Treatment and whether the test was pre-or post-initiation of the start of feeding. The interaction of treatment by time was forced into the model. Other explanatory variables (e.g. primiparous vs multiparous; breed coded as Friesian vs others; DIM at herd test) were offered to the model and included where significant (p<0.05) and/or resulted in changes in the coefficient for treatment of >20%. The estimated marginal mean and 95% confidence intervals were derived from the final model and pairwise comparison of marginal means by treatment group and time undertaken using the Bonferroni correction. Post initiation of treatment, herd tests occurred at 36, 55, 31, and 57 days after initiation of feeding for the B, W, T, and S herds, respectively.

Analyses were undertaken in STATA v16 (STATA Corp, College Station Texas, USA).

### Power analyses

*A priori* it was assumed that the bacteriological cure rate for untreated quarters would be approximately 50%, and that we wished to demonstrate a 20% absolute increase in cure (i.e. to 70%) in treatment group cows. Thus 72 infected quarters per treatment group (power = 80%, p<0.05, one-sided) were required to be available for analysis. Assuming that 70% of cows selected on the basis of having SCC >200,000 cells/mL had on average of 1.5 quarters infected per cow, and that we would lose 10% of quarter cases to follow-up (for example, for treatment of clinical mastitis, treatment with antibiotics for other reasons etc.), and we allowed for a 10% over enrolment due to clustering (i.e. quarter within cow), then we projected that we needed to screen approximately 170 cows.

## Results

The numbers of cows assessed and excluded from enrolment in the study are shown in Figure 1. Of the 267 cows initially selected based on having a SCC >200,00 cells/mL at the preceding herd test, 227 were milk sampled, with 40 not presented, diagnosed with clinical mastitis at the time of initial assessment or had very rough teat ends. Of these 227, 156 cows had one or more quarters from which bacteria were cultured, and 77 and 79 cows were assigned to the control and treatment groups, respectively. Three cows were diagnosed with clinical mastitis following initially sampling, but before commencement of feeding. These cows were excluded from the study prior to initiation of feeding.

**Figure 1.**
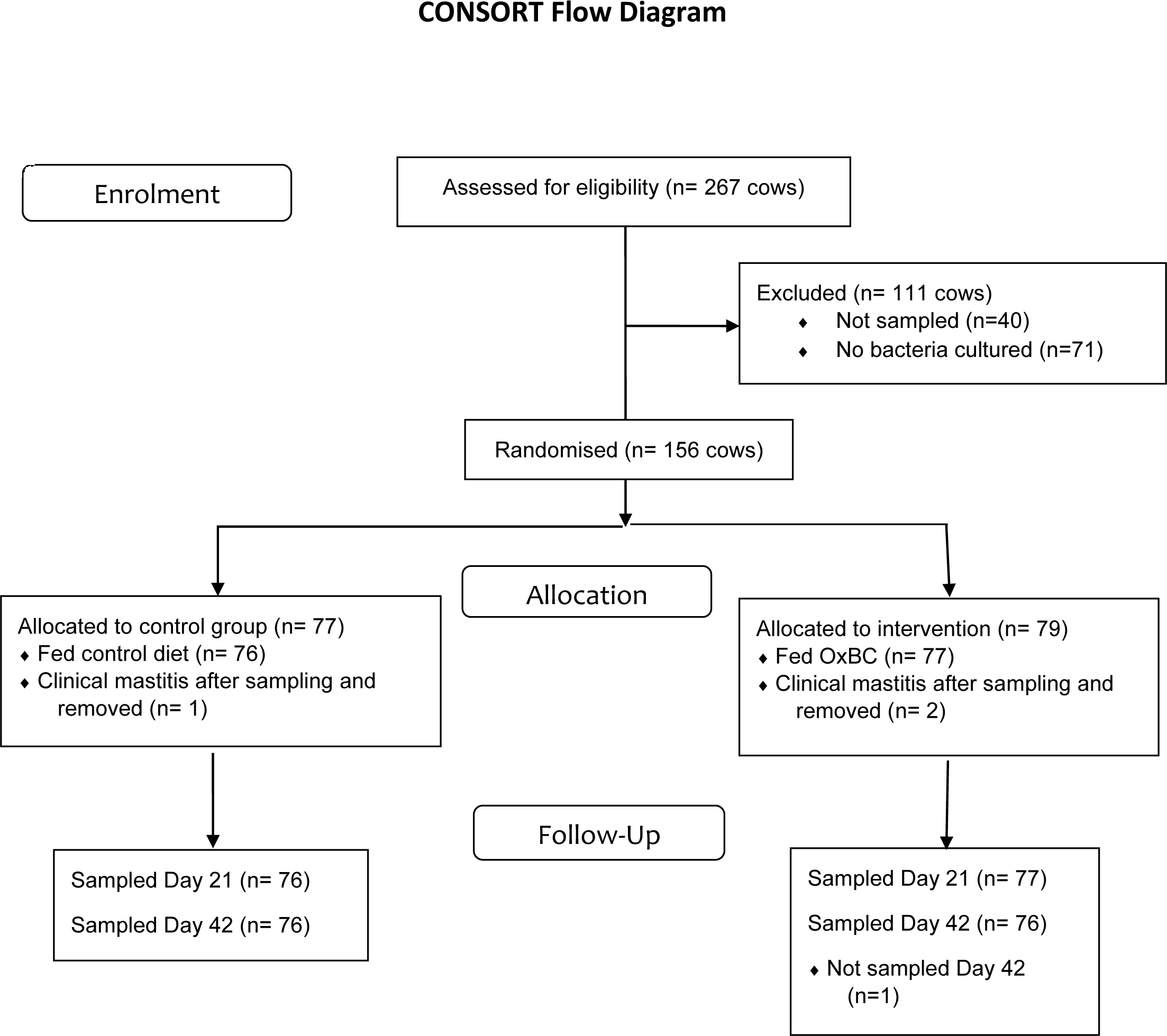
CONSORT diagram showing the numbers of dairy cows assessed, excluded and enrolled in a study to determine the effect of feeding oxidised β-carotene (OxBC) on clinical and subclinical mastitis.

Milk samples were collected from all 135 enrolled quarters in the control group on both Days 21 and 42, and from 129 and 128 quarters in the treatment group on Days 21 and 42, respectively.

One cow in the treatment group and one from the control group, both with only one enrolled quarter, were treated with systemic antimicrobials for lameness after enrolment.

There was no difference in the distribution of cows assigned to either treatment or control groups within herd, breed, age group, or DIM (all p>0.8; Table 2). The mean ln SCC recorded for enrolled cows in the herd test preceding initiation of feeding did not differ between control (6.18 (SD 0.77) cells/mL) and treatment (6.18 (SD 0.72) cells/mL) groups (p=0.96).

**Table 2.** Distribution of cows assigned to be fed a cereal-based dairy pellet that did (Treatment) or did not (Control) contain 300 mg of oxidised β-carotene (OxBC), by herd, breed, age, and days in milk at commencement of feeding.

| Variable | Control | Treatment | P-value <sup>a</sup> |
| --- | --- | --- | --- |
| Farm |  |  | 0.98 |
| B | 7 | 7 |  |
| W | 13 | 15 |  |
| T | 34 | 34 |  |
| S | 22 | 21 |  |
| Breed <sup>b</sup> |  |  | 0.95 |
| Friesian | 29 | 29 |  |
| Other | 47 | 48 |  |
| Age |  |  | 0.98 |
| Primiparous | 7 | 7 |  |
| Multiparous | 69 | 70 |  |
| Days in milk |  |  | 0.81 |
| 15–60 | 20 | 24 |  |
| 61–75 | 18 | 19 |  |
| 76–90 | 26 | 21 |  |
| >90 | 12 | 13 |  |

<sup>a</sup> Significance of difference between groups
<sup>b</sup> Where Friesian is $\geq 12/16^{\text{th}}$ Friesian

### Bacteriology and bacteriological cure

Culture results for all milk samples collected on Days 0, 21 and 24 are presented in Table 3. The most common isolates both on Day 0, and on Days 21 and 42 were CNS and *Corynebacterium* spp..

**Table 3.**
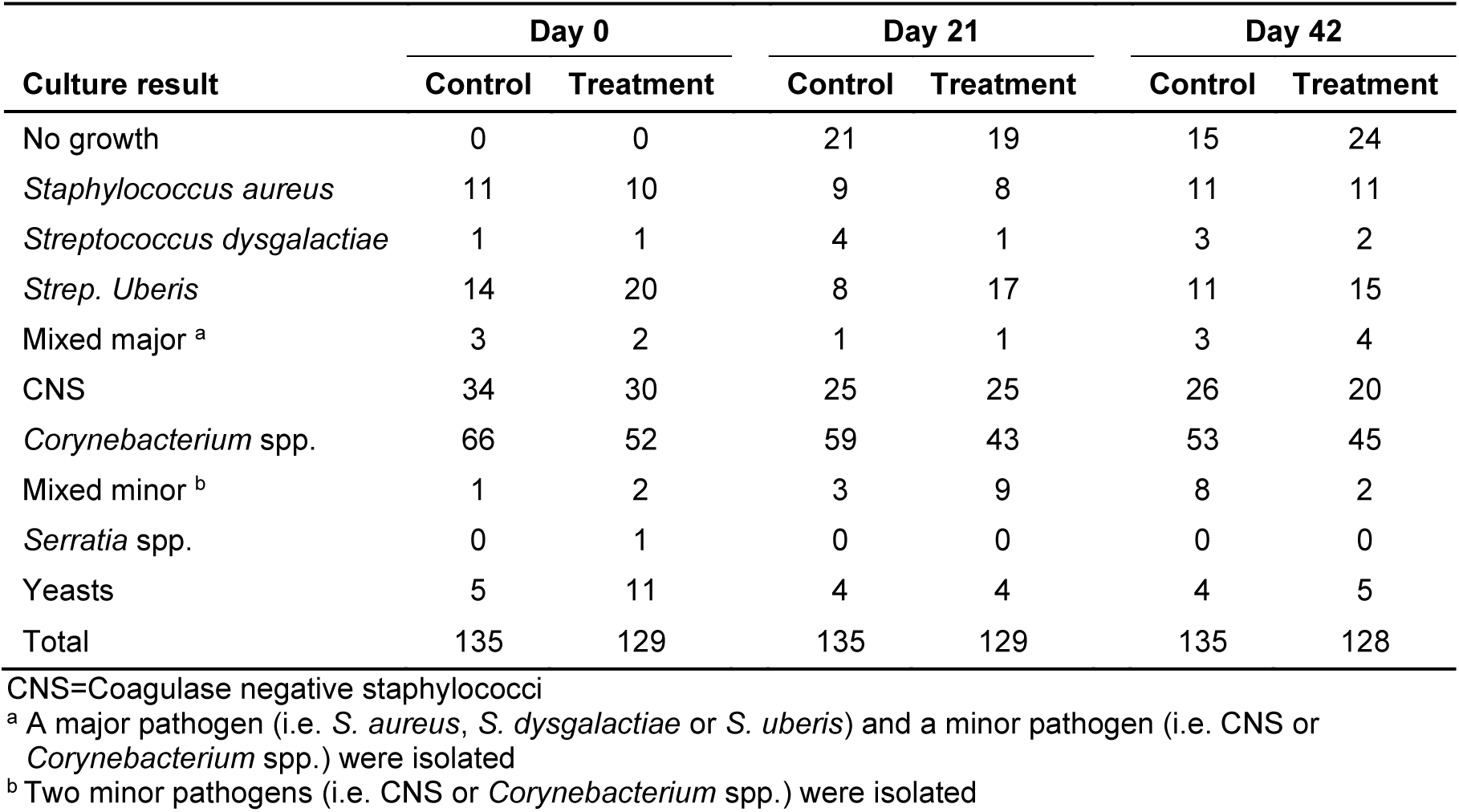
Number of milk samples collected from quarters of cows 0, 21 and 42 days after the commencement of feeding a cereal-based pellet that did (Treatment) or did not (Control) contain oxidised β-carotene, categorised by culture result.

| Culture result | Day 0 |  | Day 21 |  | Day 42 |  |
| --- | --- | --- | --- | --- | --- | --- |
|  | Control | Treatment | Control | Treatment | Control | Treatment |
| No growth | 0 | 0 | 21 | 19 | 15 | 24 |
| *Staphylococcus aureus* | 11 | 10 | 9 | 8 | 11 | 11 |
| *Streptococcus dysgalactiae* | 1 | 1 | 4 | 1 | 3 | 2 |
| *Strep. Uberis* | 14 | 20 | 8 | 17 | 11 | 15 |
| Mixed major <sup>a</sup> | 3 | 2 | 1 | 1 | 3 | 4 |
| CNS | 34 | 30 | 25 | 25 | 26 | 20 |
| *Corynebacterium* spp. | 66 | 52 | 59 | 43 | 53 | 45 |
| Mixed minor <sup>b</sup> | 1 | 2 | 3 | 9 | 8 | 2 |
| *Serratia* spp. | 0 | 1 | 0 | 0 | 0 | 0 |
| Yeasts | 5 | 11 | 4 | 4 | 4 | 5 |
| Total | 135 | 129 | 135 | 129 | 135 | 128 |

CNS=Coagulase negative staphylococci
<sup>a</sup> A major pathogen (i.e. *S. aureus*, *S. dysgalactiae* or *S. uberis*) and a minor pathogen (i.e. CNS or *Corynebacterium* spp.) were isolated
<sup>b</sup> Two minor pathogens (i.e. CNS or *Corynebacterium* spp.) were isolated

The percentage of quarters from which bacteria present on Day 0 were not isolated from samples collected on either Days 21 or 42 (bacteriological cure) was greater for cows in the treatment group (13.9 (95% CI=4.1–23.7)%) than for cows in the control group (6.9 (95% CI=4.8–9.1)%; p=0.02). The final multivariable model for bacteriological cure only included treatment group.

### New intramammary infections

More quarters developed a new intramammary infection at Day 21 or 42 for cows in the treatment group (17.9 (95% CI=6.7–29.1)%) than for quarters in cows in the control supplement group (13.0 (95% CI=4.3-21.8)%; p<0.01). The percentage of new infections was less in quarters from which a major pathogen was isolated on Day 0 (7.2 (95% CI=2.5–11.9)%) than in quarters from which a minor pathogen was isolated (17.9 (95% CI=6.4–29.4)%; p<0.01).

### Prevalence of infected quarters on Day 42

The percentage of quarters that were infected on Day 42 was less in cows in the treatment group (79.9 (95% CI=62.3–97.6)%) than the control group (88.2 (95% CI=78.4–97.9)%; p=0.009). The final multivariable model for prevalence of infected quarters on Day 42 only included treatment group.

### Quarter-level SCC

Mean quarter-level ln SCC did not differ between treatment groups (p=0.34) or Days (p=0.12), and there was no treatment by day interaction (p=0.17) (Figure 2).

**Figure 2.**
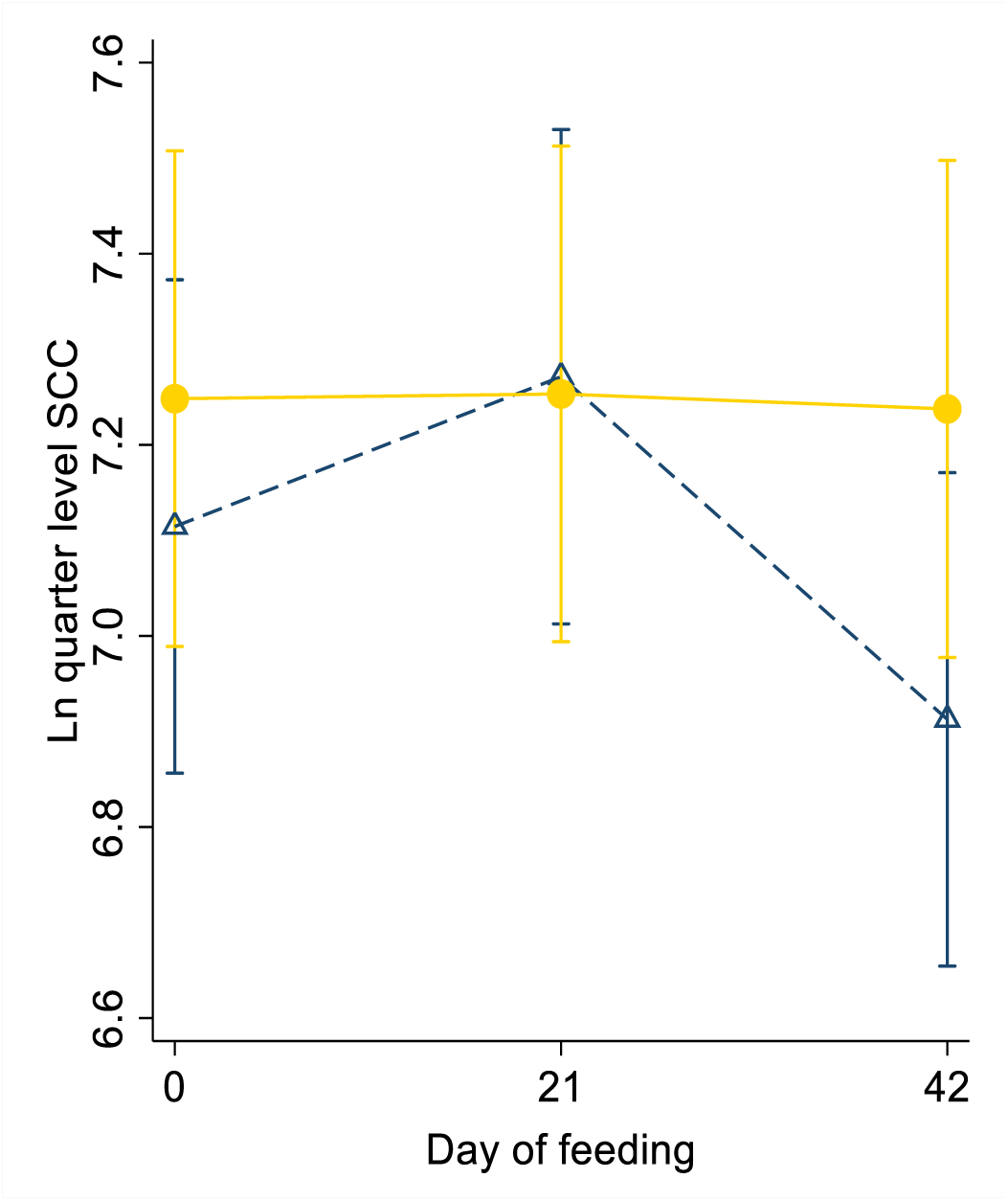
Estimated marginal mean (95% CI) ln quarter-level somatic cell counts (SCC) measured in milk samples collected from cows 0, 21 and 42 days after the commencement of feeding a cereal-based pellet that did (•) or did not (Δ) contain oxidised β-carotene.

The proportion of quarters with SCC <200,000 cells/mL was associated with a treatment by day interaction (p=0.05), with the proportion increasing between Days 21 and 42 in cows from the treatment group and decreasing in cows from the control group (Figure 3). There was no overall effect of treatment group (p=0.56) or day (p=0.64).

**Figure 3.**
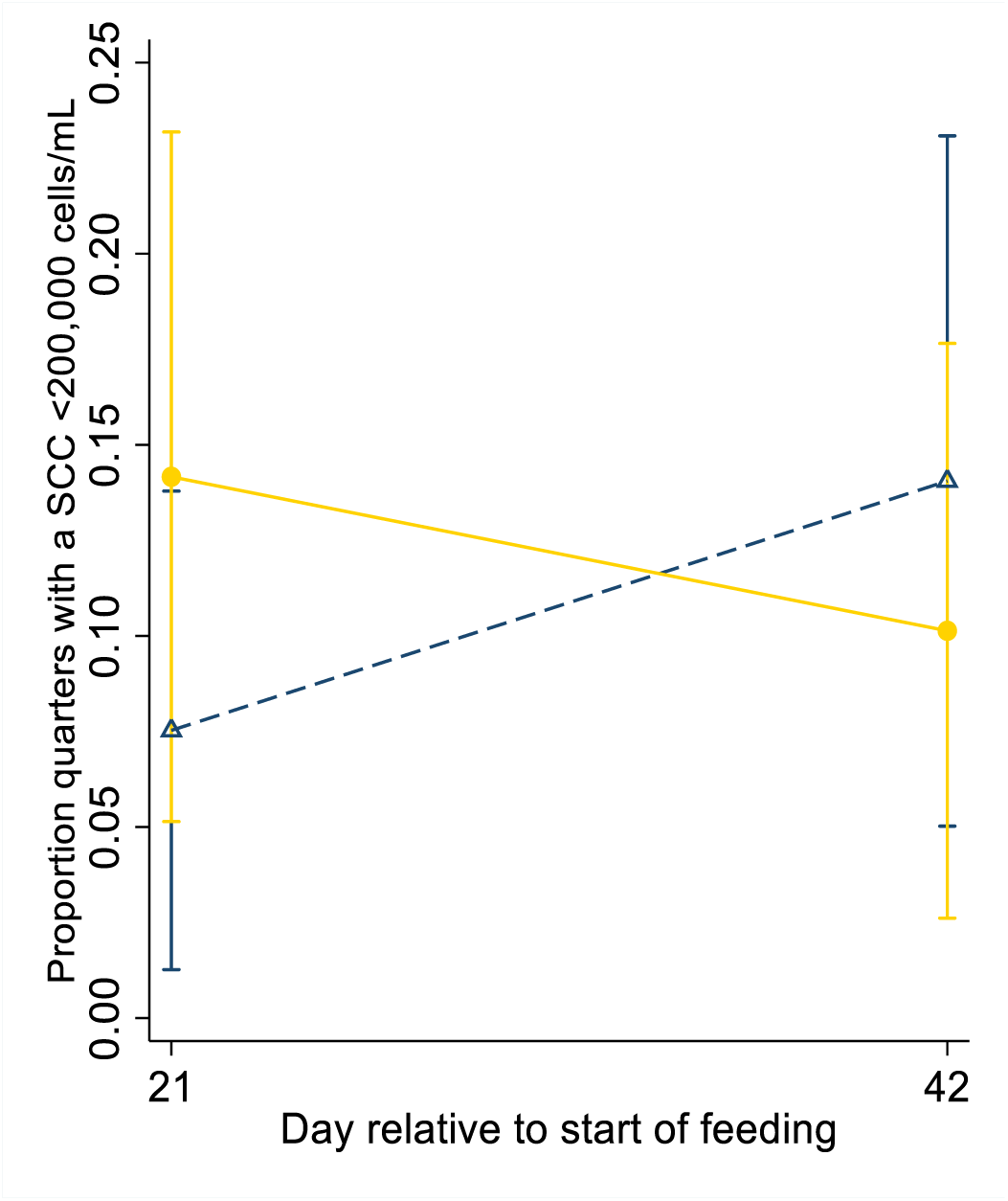
Estimated mean (95% CI) proportion of quarters with somatic cell counts (SCC) <200,000 cells/mL measured in milk samples collected from cows 21 and 42 days after the commencement of feeding a cereal-based pellet that did (•) or did not (Δ) contain oxidised β-carotene.

### Clinical mastitis incidence

Clinical mastitis was diagnosed between Days 0 and 42 in four cows and six quarters from the control group and one cow and one quarter from the treatment group. The incidence of quarters diagnosed with clinical mastitis was lower in cows from the treatment group (1/129 (0.78 (exact binomial 95% CI=0.02–4.24)%) than in cows from the control group (6/135 (4.44 (exact binomial 95%CI=1.65–9.42)%); OR=0.17 (95% CI=0.03-0.82); p=0.03). The one quarter of the one cow in the treatment group with clinical mastitis was diagnosed on Day 41. This cow was chronically infected with *S. aureus*.

### Cow-level SCC, volume, fat percentage and protein percentage at subsequent herd test

There was no difference between treatment and control groups in cow-level SCC, milk yield or milk composition recorded at the herd test following the start of the study period (Supplementary Figures 3-6).

## Discussion

This controlled randomised intervention study demonstrated that oral supplementation of lactating dairy cows with subclinical intramammary infections with an oxidised β-carotene product resulted in an increased bacteriological cure, a reduced prevalence of intramammary infection on Day 42, and a reduced incidence of clinical mastitis by Day 42, relative to control cows. Supplementation had no effect on quarter-level SCC or on milk yield or composition.

A total of 135 and 129 quarters were allocated to the treatment and control group, respectively. *A priori* 72 quarters per group were calculated as being required. However, the initial power analysis assumed that 50% of the control group quarters would spontaneously cure and that we wished to demonstrate a 20% increase in the cure rate with treatment. The bacteriological cure rate was only 7% in the control group and 14% in the treatment group. The initial assumption that there would be a 50% cure in the control group was based on an earlier preliminary New Zealand study where more than one third of infected but untreated quarters underwent spontaneous cure (Douglas *et al.* 1997). However, in that study the inclusion criteria did not include quarter-level SCC. Thus, the more stringent enrolment criteria in the current study may have resulted in more significant infections being selected and hence a lower spontaneous cure rate. Despite the smaller difference in cure rate between the treatment and control quarters than expected, the over-enrolment (relative to initial study design) resulted in sufficient statistical power to demonstrate a difference in bacteriological cure rate between treatment groups.

The current study results support and extend previous studies that have demonstrated positive effects on immune function and milk quality of vitamin A or β-carotene supplementation of lactating dairy cows. It should be noted that while OxBC is derived from β-carotene, it does not contain intact β-carotene, vitamin A or the ability to activate the retinoic acid receptor system (Burton *et al.* 2014; Johnston *et al.* 2014). Instead OxBC is predominantly composed of beta-carotene-oxygen copolymers that a form spontaneously when beta-carotene is exposed to oxygen in air (Burton *et al.* 2014). Previous reports indicate that OxBC has the ability to increase the level of cellular TLR-2 and −4 along with their co-receptor CD14, as well as to enhance the activity of downstream innate immune effector mechanisms responsible for clearing bacterial infections (Burton *et al.* 2014; Johnston *et al.* 2014). The absence of intact beta-carotene as well as any vitamin A associated activity for OxBC has led some authors to propose that the copolymers present within OxBC are the actual agents responsible for many of the non-provitamin A actions of beta-carotene and other carotenoids (Johnston *et al.* 2014). Up-regulation of TLR-2, TLR-4 and CD14 coupled with modulation of chemokine levels and immune effector mechanisms have been shown to improve pathogen recognition and bacterial clearance from the mammary gland (De Schepper *et al*. 2008; Jin *et al*. 2014). While immunological parameters were not evaluated in the present study, the previously reported effects of OxBC on pattern recognition receptor and chemokine levels coupled with enhanced macrophage activity offer plausible mechanisms to explain the observed increase in bacteriological cure rate in the OxBC treated groups. The results of studies with OxBC in other species are consistent with the observed increased cure rate in dairy cattle. More specifically, dietary supplementation with OxBC led to significantly lower levels of pathogen in the gut and improved growth performance of *Clostridium perfringens-*challenged broiler chickens (Kang *et al*. 2018). Also, inclusion of OxBC in the diets of gestating and lactating sows led to a significant increase in immunoglobulin concentrations in both the colostrum and milk (Chen *et al*. 2020).

Although feeding of OxBC resulted in a lower proportion of quarters remaining infected at Day 42, 80% of infections persisted at Day 42. As identified above, the spontaneous cure rate of untreated subclinical intramammary infections was previously reported to be 33% (Douglas *et al.* 1997). In contrast, the bacteriological cure rate in quarters in the control group was only 7%, and only 12% of quarters were uninfected at Day 42 in the current study. One implication of these findings is that assumptions that subclinical intramammary infections spontaneously resolve and hence may have limited longer term impact on cows needs to be reviewed. Further longitudinal studies under New Zealand management conditions are required to assess the duration of these subclinical infections, and secondly to define the impact of such infections on milk yield, risk of transfer infection to other cows within the herd, and on risk of culling.

Paradoxically, quarters from cows in the treatment group had a higher new intramammary infection rate over the 42 days compared with cows in the control group, yet at the final sampling at Day 42 fewer quarters in the treatment group remained infected. One potential explanation for this is that the presence of a major pathogen intramammary infection at Day 0 was associated with a reduced rate of new intramammary infections over the subsequent 42 days, compared with presence of a minor pathogen. As OxBC feeding resulted in a higher cure rate of infections, including major pathogen infections, this potentially may have increased the susceptibility of these quarters to reinfection. However, the net effect of cure and of reinfection was that the prevalence of infection at Day 42 was lower in the treatment than control group.

Feeding of OxBC resulted in an 83% reduced odds of clinical mastitis during the 6-week period of the study. This may have been a result of the reduced prevalence of intramammary infection and hence a lower likelihood of clinical mastitis occurring but may also reflect modulation of the inflammatory response resulting in reduced severity of clinical signs amongst cows with intramammary infection. In a calf respiratory disease model, calves fed OxBC had less severe clinical signs (Duquette *et al.* 2014). Further evidence of the ability of OxBC to modulate the inflammatory response is provided by a study with gestating/lactating sows which showed that animals receiving dietary OxBC had significantly lower TNFa and IL-8 levels in colostrum compared to controls. The reduction in TNFa concentration was also evident in milk samples taken at day 14 of lactation (Chen *et al*. 2020).

Despite a higher bacteriological cure rate in the quarters from cows fed OxBC in the current study, there was no difference in quarter or cow level SCC between treatment groups. This may be due to the relatively low cure rate and any effect on SCC being masked by the presence of the remaining bacteria. It is also interesting to note that less than 20% of cured quarters had SCC ≤200,000 cells/ml by Day 42, illustrating the long period of inflammation that occurs in the mammary gland.

No effect of OxBC treatment on milk yield or milk composition was detected. However, the study was not designed to look for such effects, and given the sample size, very substantial differences in milk yield and composition would need to have occurred for it to be detectable within the current study.

It is beyond the scope of the present study to assess the economic benefit of feeding OxBC. However, the reduced prevalence of infection at Day 42 and reduced incidence of clinical mastitis may result in reduced cow to cow spread of pathogens, and reduced costs associated with antimicrobial treatments and milk discard. The bacteriological cure rate resulting from feeding OxBC may be lower than if these quarters had been treated with antimicrobials, but the cost of feeding OxBC should be markedly lower than using antimicrobials due to there being no milk discard. In addition, there is no increased risk of development of antimicrobial resistance.

In conclusion, oral supplementation of lactating dairy cows with subclinical intramammary infections with OxBC resulted in a higher bacteriological cure rate, a lower prevalence of intramammary infection at 42 days, and a lower incidence of clinical mastitis than in cows not fed OxBC. There was no effect of feeding OxBC on quarter-level SCC, or on milk yield or composition.

## Acknowledgements

The assistance of the herdowners and staff in undertaking this study is acknowledged.

The field and laboratory staff of Cognosco including Elizabeth Blythe, Cathy Yanez, Bev Brownlie, Yvette Macpherson, and Ali Karkaba are thanked for their excellent work on this study which involved daily trips to farms for nearly 3 months as well as a requirement for rapid turnaround of laboratory results.

Avivagen funded this study.

*Non peer reviewed

## References

Burton GW, Daroszewski J, Nickerson JG, Johnston JB, Mogg TJ, Nikiforov GB. β-carotene autoxidation: Oxygen copolymerization, non-vitamin a products, and immunological activity. Canadian Journal of Chemistry 92, 305–16, 2014

Chen J, Zhang Y, Lv Y, Tian M, Cheng L, Chen F, Zhang S, Guan W. Effects of dietary fully oxidized β-carotene polymers during late gestation and lactation on productivity and sow immune status. British Journal of Nutrition. 10.1017/S0007114520002652

Chew BP, Hollen LL, Hillers JK, Herlugson ML. Relationship between vitamin A and β-carotene in blood plasma and milk and mastitis in Holsteins. Journal of Dairy Science 65, 2111–8, 1982

Chew BP, Park JS. Carotenoid action on the immune response. The Journal of Nutrition 134, 257S–61S, 2004

De Schepper S, De Ketelaere A, Bannerman DD, Paape MJ, Peelman L, Burvenich C. The toll-like receptor-4 (tlr-4) pathway and its possible role in the pathogenesis of *Escherichia coli* mastitis in dairy cattle. Veterinary Research 39, 5, 2008

*Douglas VL, Holmes CW, Williamson NB, Steffert IJ. Use of individual cow somatic cell counts, electrical conductivity, and the rapid mastitis test on individual quarters to diagnose subclinical mastitis in ealry lactation, with an economic assessment of antibiotic therapy. Proceedings of the 14th Annual Seminar of the Society of Dairy Cattle Veterinarians of the New Zealand Veterinary Association 80–90, 1997

Duquette SC, Fischer CD, Feener TD, Muench GP, Morck DW, Barreda DR, Nickerson JG, Buret AG. Anti-inflammatory effects of retinoids and carotenoid derivatives on caspase-3-dependent apoptosis and efferocytosis of bovine neutrophils. American Journal of Veterinary Research 75, 1064–75, 2014

Halasa T, Huijps K, Osterás O, Hogeveen H. Economic effects of bovine mastitis and mastitis management: A review. Veterinary Quarterly 29, 18–31, 2007

*Hogan JS, Gonzalez RN, Harmon RJ, Nickerson SC, Oliver SP, Pankey JW, Smith KL. Laboratory Handbook on Bovine Mastitis. National Mastitis Council Inc., Madison, WI, USA, 1999

Jin L, Yan S, Shi B, Bao H, Gong J, Guo X, Li J. Effects of vitamin a on the milk performance, antioxidant functions and immune functions of dairy cows. Animal Feed Science and Technology 192, 15–23, 2014

Johnston LA, Chew BP. Peripartum changes of plasma and milk vitamin A and *β*-carotene among dairy cows with or without mastitis. Journal of Dairy Science, 67, 1832–40, 1984

Johnston JB, Nickerson JG, Daroszewski J, Mogg TJ, Burton GW. Biologically active polymers from spontaneous carotenoid oxidation: A new frontier in carotenoid activity. PLOS ONE 9, e111346–e, 2014

Kang M, Oh J-Y, Cha S-Y, Kim W-I, Cho H-S, Jang H-K. Efficacy of polymers from spontaneous carotenoid oxidation in reducing necrotic enteritis in broilers. Poultry Science 97(9), 3058–3062, 2018

*Mein GA, Neijenhuis F, Morgan WF, Reinemann DJ, Hillerton JE, Baines JR, Ohnstad I, Rasmussen MD, Timms L, Britt JS, Farnsworth R, Cook N, Hemlin T. Evaluation of bovine teat conditions in commercial dairy herds 1. Non-infectious factors. In: Proceedings of the 2nd International Symposium on Mastitis and Milk Quality 347–51, 2001

Morris RS. The depression of quarter milk yield caused by bovine mastitis, and the response of yield to successful therapy. Australian Veterinary Journal 49, 153–6, 1973

Mwakipesile SM, Holmes CW, Moore YF. Antibiotic therapy for subclinical mastitis in early lactation; effects on infection, somatic cell count and milk production. New Zealand Veterinary Journal 31, 192–5, 1983

Oliveira RC, Guerreiro BM, Morais Junior NN, Araujo RL, Pereira RAN, Pereira MN. Supplementation of prepartum dairy cows with β-carotene. Journal of Dairy Science 98, 6304–14, 2015

*Scherf H, Frye TM, Williams SN. Vitamin A and β-carotene: A nutritional approach to the control of mastitis in dairy cattle. Proceedings of the National Mastitis Council 22, 77, 1994

Sordillo LM, Shafer-Weaver K, DeRosa D. Immunobiology of the mammary-gland. Journal of Dairy Science 80, 1851–65, 1997

Swinkels JM, Hogeveen H, Zadoks RN. A partial budget model to estimate economic benefits of lactational treatment of subclinical *Staphylococcus aureus* mastitis. Journal of Dairy Science 88, 4273–87, 2005

Tjoelker LW, Chew BP, Tanaka TS, Daniel LR. Effect of dietary vitamin A and β-carotene on polymorphonuclear leukocyte and lymphocyte function in dairy cows during the early dry period. Journal of Dairy Science 73, 1017–22, 1990

